# Characterization of *ATM* gene expression and evaluation of Reactive Oxygen Species in Silibinin-treated SKBR3 cells

**DOI:** 10.64898/2026.07.02.736131

**Authors:** Negar Sadat Nademi, Nasrin Motamed

## Abstract

**Background:** Reactive Oxygen Species (ROS) are the small, unstable and highly reactive species, having DNA oxidizing ability. Oxidation of the DNA’s purine and pyrimidine bases can lead to single or double strands in this macromolecule. In this situation, the ATM molecule, a serine-threonine kinase, targets several proteins for phosphorylation, which causes the cell cycle to stop and the DNA damage repair begins. It has previously been proven that natural polyphenols have the cancer inhibiting properties due to their high efficacy and low side effects. Silibinin is the main herbal and medical ingredient in Milk Thistle (*Silybum marianum*) is a polyphenol flavonolignan, which has been widely considered as an antioxidant and anticancer agent. The purpose of the present study was to investigate the *ATM* gene expression and measurement of reactive oxygen species (ROS) in SKBR3 cell line, treated with Silibinin.

**Materials and Methods:** At first, the SKBR3 cell line was cultured in RPMI1640 culture medium and MTT assay was carried out to evaluate the Silibinin cytotoxicity. Flow Cytometry was carried out for cell cycle analysis, apoptotic induction, and ROS detection. While, Real Time PCR was used to evaluate the *ATM* gene expression in the Silibinin-treated and un-treated SKBR3 cells.

**Results:** Present results have shown that 150 µ*M* Silibinin had the most significant cytotoxicity and apoptotic induction influence after the treatment period of 48 h. Flow cytometry data have shown that Silibinin induced considerable amount of apoptosis and caused cell cycle arrest at G1/S phase and induced production of ROS. Real-time PCR results have revealed that Silibinin increased the *ATM* expression in SKBR3 cell line.

**Conclusion:** Silibinin causes increased *ATM* gene expression by inducing ROS production, which initiates cell cycle arrest and apoptotic induction in SKBR3 cells line.

## Introduction

According to the World Health Organization’s recent reports, 14 million individuals get affected with cancer, with 8.2 million mortalities. It is estimated that in the next two decades, the cancer incidence would reach 22 million worldwide [1]. According to the World Health Organization (WHO), the incidence of cancer in Iran would be about 131,191 new cases and 79,136 deaths among its population of nearly 84 million in 2020 [41]. More recent global estimates confirm that this burden has continued to rise: breast cancer alone accounted for approximately 2.3 million new cases and 666,000 deaths worldwide in 2022 [34].

Cells become cancerous by undergoing unusual and abnormal changes in apoptotic and cell cycle pathways, causing metabolic disorders and genetic instabilities [5]. Apoptosis is a programmed cell death that works in parallel and opposite to the cell proliferation. Both of these processes occur in response to some stimuli and are also controlled by the process of morphological differentiation. In human body, an average of 60 billion cells die every day with scheduled cellular death [6, 7]. Normal amount of apoptotic process exerts a control on tissue proliferation and growth, while retarded apoptotic pathway results in the emergence of cancer cells. Many factors are involved in the apoptotic process. Pro-apoptotic factors have the ability to initiate apoptosis, while anti-apoptotic processes work by inhibiting apoptotic pathways [8]. The expression and activity of these factors varies depending on the type of cell and stimulus [9].

Oxygen is an important element in the process of cellular respiration and acts as the final electron receptor in the ATP synthesis metabolism [10, 11]. Sometimes the oxygen reacts with the electrons, expelled from the electron transport chain in mitochondrial membranes and results in the production of free radicals, that are called reactive oxygen species (ROS) [11, 12]. ROS have the potential to react with different molecules in the mitochondria.

Whenever the intracellular factors are unable to combat the ROS, cellular proteins, DNA and lipid moieties suffer from oxidative damages [13, 14]. The oxidation of DNA’s pyridine and purine bases can lead to single or double strand breaks in this macromolecule.

In this situation, the ATM molecule, a serine-threonine kinase, targets several proteins for phosphorylation, which causes cell cycle arrest and DNA repair [15, 16]. P53 protein is one of the most important targets of ATM, which in turn results in the expression of BAX, GADD45, p21 and PUMA proteins that control cell cycle progression, DNA damage repair and apoptotic induction. A number of edible herbs have anti-oxidant properties and can play an important role in preventing and treating cancer [17]. One of these herbs is the milk thistle (*Silybum marianum*), the seed extract of which is rich in polyphenolic flavonoids. Anti-oxidant compound Silibinin constitutes about 90% of the milk thistle extract [18, 19]. Silibinin has been proven to exert anticancer effects against breast, colon, lung, liver, bladder and prostate cancers [18, 20-22].

So, the present study was designed to evaluate the ROS production and *ATM* gene expression in the SKBR3 cell line, treated with different concentrations of Silibinin.

## Materials and Methods

### Chemicals and Cell Line

The SKBR3 cell line of human breast cancer adenocarcinoma, used in this study, was obtained from the National Cell Bank of Iran, Pasteur Institute, Tehran, Iran. SKBR3 cell line was cultured in RPMI1640 culture medium (Invitrogen, Carlsbad, CA, USA) with 15% Fetal Bovine Serum (FBS)and Penicillin/Streptomycin (1 ml/100ml culture medium) antibiotic, and incubated at 37 °C and 5% CO_2_ for 24 h. Silibinin was purchased from Merck, Darmstadt, Germany, and dissolved in 1% sterile DMSO and at 20 °C for further use.

### Cell culture and MTT Assay

In order to evaluate the Silibinin cytotoxicity at the concentrations of 50 to 200 μm and obtaining its IC50 (concentration of a drug that inhibits 50% cell growth) dosage, the SKBR3 cells were treated with it for 24, 48 and 72 hand the obtained results were compared with those of the control group. After the cells reached 90% confluency, the trypan blue staining was carried out and the cells were then counted and seeded at the number of 5000 cells per well of 96-well plates and incubated for 24 h. Then, the cells were treated with various concentrations of Silibinin and underwent final incubation for the treatment periods of 24, 48 and 72 h.

MTT assay kit was used to evaluate cytotoxic effects of Silibinin as per manufacturer’s instructions. Cells were incubated for 4 h with the MTT reagent and the formazan crystals were formed as a result of MTT’s reduction with the help of active succinate dehydrogenase enzyme in the mitochondria of live cells. The purple formazan crystals were solubilized after addition of organic solvent, DMSO, prior to colorimetric analysis at the wavelength of 570 nm via the ELISA reader. Percentage of vital capacity (survival) of SKBR3 cells was obtained using following relation:

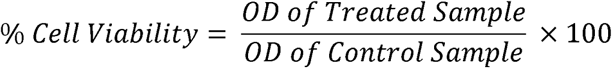

### IC50 Determination

In order to determine IC50 dose of Silibinin against SKBR3 cells, the average of three absorbance readings pf each sample was considered and calculated with the help of Pharm Pharmacological Calculating System (PCS) statistical package (Springer Verlag, USA) and IC50 values of each Silibinin concentration were determined after 24, 48 and 72 h. The corresponding graphs were drawn using SPSS (Ver. 22.0) software.

### Apoptotic induction analysis by Flow Cytometry

The apoptotic induction capacity of different doses of Silibinin was measured by flow cytometry, using propidium iodide (PI) and annexin V-FITC apoptosis kit (Bioscience, USA). As live and metabolically active cells can propel the PI out of the cell membrane territory, the PI-positive cells are considered dead. While, annexin V was used to determine cell death by flow cytometry. For this, 750,000 cells were planted in a filtered flask and incubated for 48 h. the cells were transferred to the separate falcons without passaging, and 1.5 ml trypsin-EDTA was added and the cells were centrifuged at 1500×g. the supernatant was discarded and the cells were washed twice with cold PBS and then the binding buffer was added to the cells after discarding PBS. The annexin V was added to the resultant cells at the concentration of 5 µl/100 µl cells and the samples were incubated at room temperature for 20 min, then the PI was finally added to them and the saples were transferred to the flow cytometric tubes for analysis. The results were analyzed by ANOVA test using SPSS 22.0 software.

### Cell cycle analysis

The SKBR3 cells were cultured at the 80% confluency and the FBS was removed from them in order to achieve uniform cell cycle conditions. Then, the cells were trypsinized and washed twice with PBS and the resultant sediments were slowly pipetted into Pyrrolo-benzodiazepine (PBD), in order to study the cell cycle arrest. About 900 μl of 70% ethanol was added to the cell suspension, centrifuged and the supernatant was discarded following washing of sediments with PBS. Washing with PBS and the centrifugation process was carried out twice. The PBS was slowly removed and 500 μl of PI Master Mix was added and the falcons were placed in an aluminum foil at 37 °C for 30 min before flow cytometric analysis.

### ROS detection by DCFDA Assay

2’,7’ –dichlorofluorescin diacetate (DCFDA) - Cellular Reactive Oxygen Species Detection Assay Kit (Kiazist, Iran) was used to identify reactive species, resulted by the cytotoxic effect of Silibinin on SKBR3 cells. DCFDA (also known as H2DCFDA) is a fluorogenic dye that measures hydroxyl, peroxyl and other ROS activity within the cell. After diffusion into the cell, DCFDA/H2DCFDA is deacetylated by cellular esterases to a non-fluorescent compound, which is later oxidized by ROS into 2’,7’–dichlorofluorescein (DCF). DCF is a highly fluorescent compound, which can be detected by fluorescence spectroscopy with excitation/emission at 495 nm/529 nm. The analysis was carried out according to the manufacturer’s instructions.

### Real Time RT-PCR analysis

The cells were incubated in RPMI1640 culture medium and treated with 150 μM Silibinin for 48 h. The cells were then harvested in order to extract total cellular RNA using TRIzol method kit as per manufacturer’s protocol. The total RNA quantity was measured with ND-1000 Nanodrop spectrophotometer (Nanodrop Technologies Inc.) The RevertAid™ First (Fermentas) reverse transcription kit reaction was used for real time RT-PCR. The final volume of each reaction mixture was 20 μl, in which 1000 ng total RNA was present for RT process and to be converted into cDNA.

At first, each RNA sample was taken at 1000 ng quantity (depending on the RNA concentration, the volume may vary) in the 0.2 ml vials with UltraPure DEPC (Diethyl pyrocarbonate) - Treated Water, at a final volume of 10 μl for each reaction. One microliter of Random Hexamer Primer was added and placed in thermocycler at 65 °C for 5 min. To make Master Mix, 1 μl of dNTP, 1 μl of Reverse Transcriptase and 1 μl of RiboLock RNase Inhibitor, 4 μl of Reverse Transcriptase Buffer (5X) and 2 μl of DEPC water were added for each reaction, and run in thermocycler for 5 min at 25 °C, 60 min at 42 °C and 10 °C at 70 °C. The synthesized cDNA was stored at -20 °C.

### Investigation of ATM expression

To investigate the effect of Silibinin on ATM gene expression in SKBR3 cell lines in control and Silibinin-treated samples, Real Time PCR was performed using Rotor-Gene 6000. The final volume of each reaction mixture was 10 μl and constituted Maxima SYBR green master mix, 2 μl cDNA product, 0.5 μl ATM, GAPDH primers and 7 μl of nuclease-free water. The applied temperature conditions included an initial activation step at 95 °C for 30 sec, 45 cycles of 5 seconds at 95 °C for denaturation and finally the annealing / extension stage. To calculate the accuracy and specificity of PCR product, melting curve was drawn. Finally, for the calculation of the relative mRNA copy number, 2ΔΔCT formula was used. The promer sequences are given in Table 1.

### Statistical analysis

SPSS 22.0 software was used for statistical analysis. The statistical difference between the control and treated groups was determined using Paired Student’s *t*-test.

## Results

### MTT cell viability assay in SKBR3 cells

CKBR3 viability analysis was done on the SKBR3 cells at 50 – 200 µM/l concentrations of Silibinin after the culture periods of 24, 48 and 72 h. The cell viability was then assayed by MTT analysis. Percent cell viability rate of Silibinin-treated SKBR3 cells is shown in **Figure 1**.

**Figure 1.**
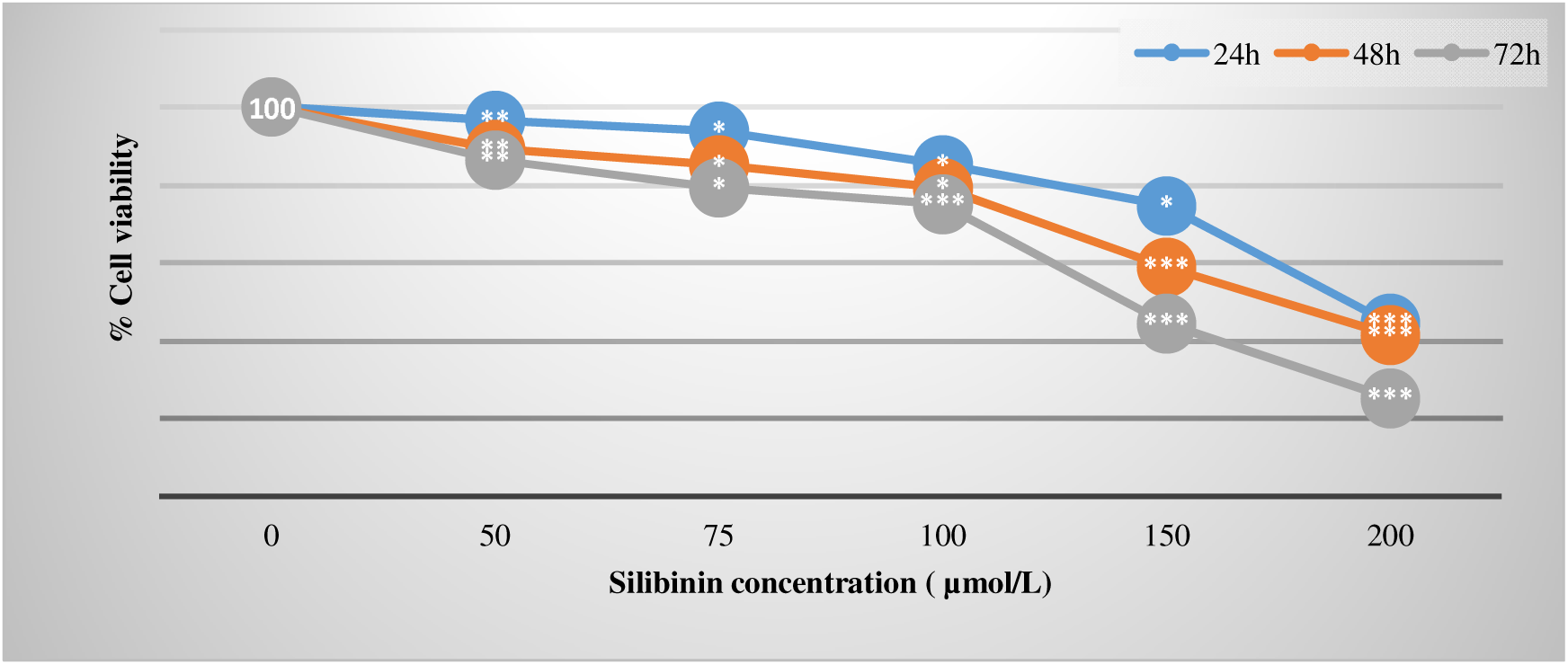
MTT assay of SKBR3 cell viability following treatment with different concentrations of Silibinin for 24, 48, and 72 h. Data are presented as mean ± SD. Cell viability was calculated relative to the untreated control group (0 µmol/L), which was set as 100%. *P < 0.05, **P < 0.01, ***P < 0.001 versus control.

According to present results, the Silibinin exerted the cytotoxicity in SKBR3 cells, which is the function of concentration and time. In this way, by increasing the concentration of Silibinin, the cell viability percentage reduces, and this decrease was prolonged by prolonging the exposure time of the cells upto 48 and 72 h. the results were significant as compared with the control group. All data, obtained from MTT assay, were analyzed using Microsoft Excel software. The IC50 dose of the Silibinin was also determined during this experiment.

### IC50 analysis

**Figure 2** shows the IC50 results of SKBR3 cells, treated with different concentrations of Silibinin during the incubation periods of 24, 48 and 72 h. As shown in **Figure 2**, the IC50 value of Silibinin against SKBR3 cells decreased in a clear time-dependent manner, from 234.49 µM at 24 h to 161.4 µM at 48 h and 133.72 µM at 72 h, confirming that prolonged exposure enhanced the cytotoxic potency of Silibinin in this cell line.

**Figure 2:**
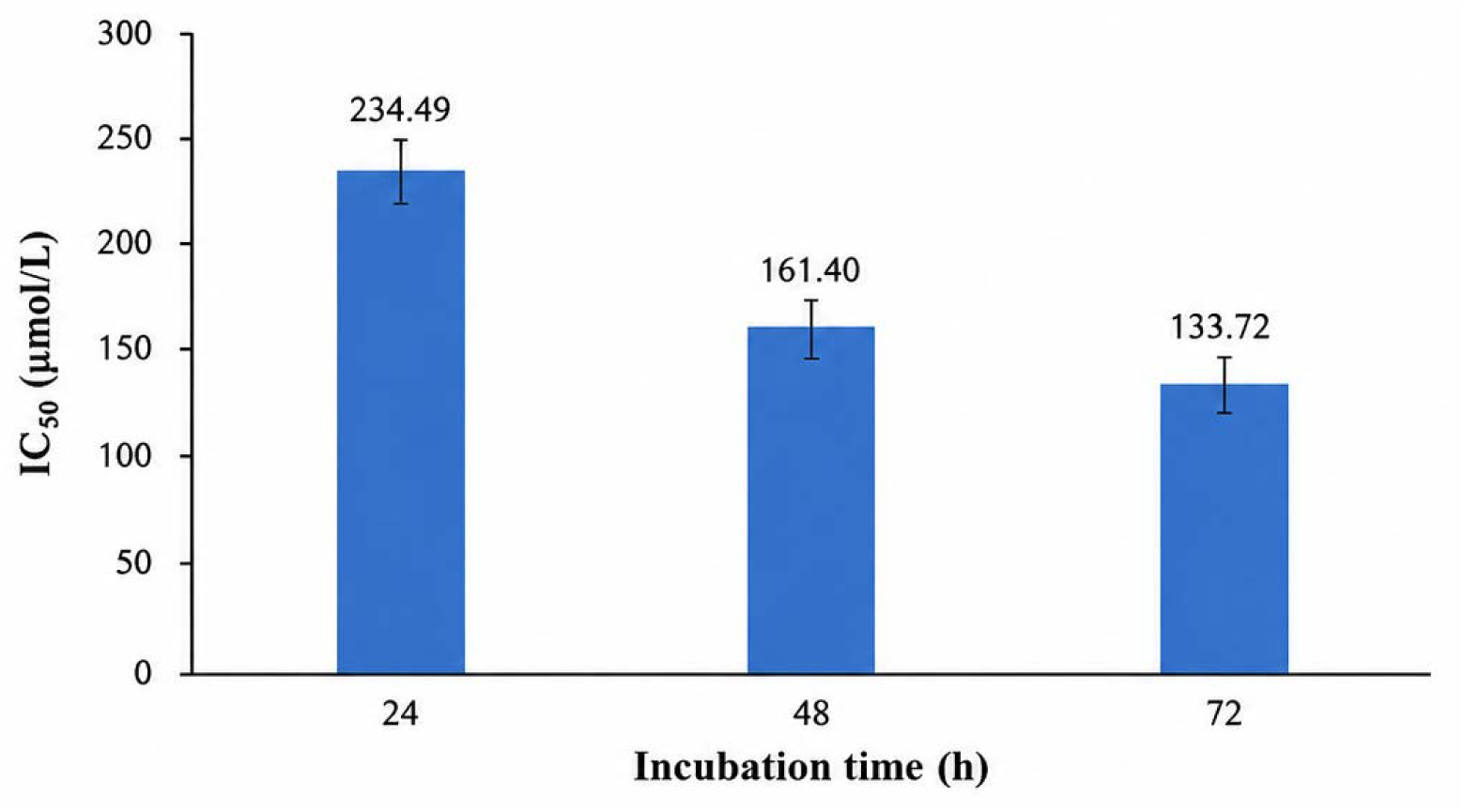
Comparison of ICll values of Silibinin in SKBR3 cells following 24, 48, and 72 h of treatment. ICll values were determined from MTT dose–response curves. Data are presented as mean ± SD from three independent experiments (n = 3).

This trend is illustrated graphically below, together with the concentration-response curves used to derive each IC50 value.

### Apoptotic analysis in SKBR3 cells by Flow Cytometry

Phosphatidylserine (PS) externalization is an early hallmark of apoptosis. During apoptosis, PS translocates from the inner to the outer leaflet of the plasma membrane, where it is detected by Annexin V. Propidium iodide (PI) enters cells with compromised membrane integrity and is used to distinguish late apoptotic or necrotic cells. As shown in **Figure 3**, treatment of SKBR3 cells with 150 µM Silibinin increased the proportion of apoptotic cells compared with the untreated control. In the control group, approximately 78.0% of cells were Annexin V−/PI− (Q4, viable cells), whereas this proportion decreased to 73.0% following Silibinin treatment. The percentage of early apoptotic cells (Annexin V+/PI−, Q3) increased from 7.7% in the control group to 23.3% in the treated group. In contrast, the proportion of late apoptotic cells (Annexin V+/PI+, Q2) remained low, changing from 1.5% in the control group to 0.2% in the treated group.

**Figure 3.**
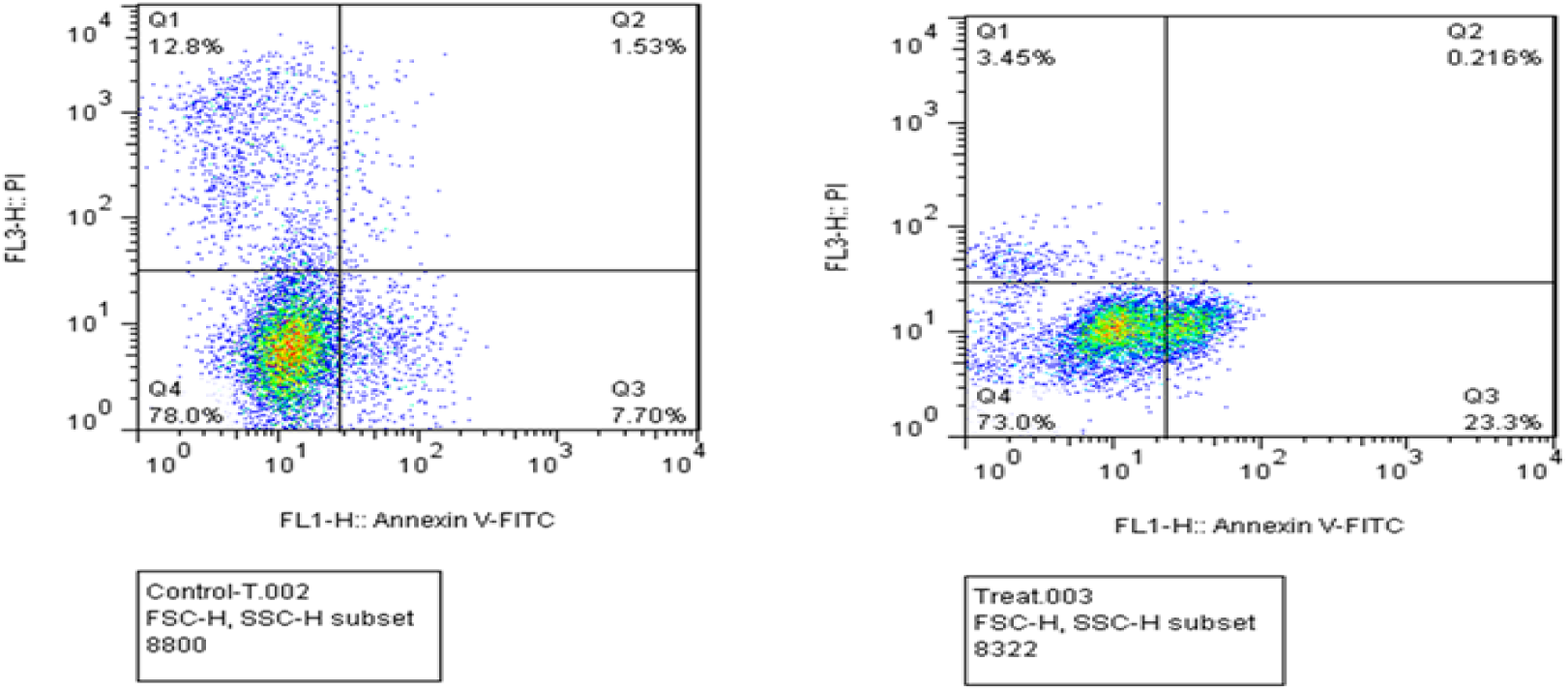
Representative Annexin V–FITC/propidium iodide (PI) flow cytometric analysis of apoptosis in SKBR3 cells following treatment with 150 µM Silibinin. The left panel shows untreated control cells, and the right panel shows Silibinin-treated cells. Q4 (Annexin V−/PI−) represents viable cells, Q3 (Annexin V+/PI−) early apoptotic cells, Q2 (Annexin V+/PI+) late apoptotic/secondary necrotic cells, and Q1 (Annexin V−/PI+) necrotic cells. Silibinin treatment increased the proportion of early apoptotic cells while reducing the percentage of viable cells compared with the untreated control.

### Cell cycle analysis in SKBR3 cells via Flow Cytometry

**Figure 4** shows the Flow cytometry graphs of cell cycle in untreated and Silibinin- treated SKBR3 cells. According to the results, the number of cells at the G1 stage is more in the Silibinin- treated cells as compared to the control group. While, a very small percentage of SKBR3 cells can be seen in G0, G1, S, and G2/M stages of cell cycle in the treated samples, as compared to the control group.

**Figure 4:**
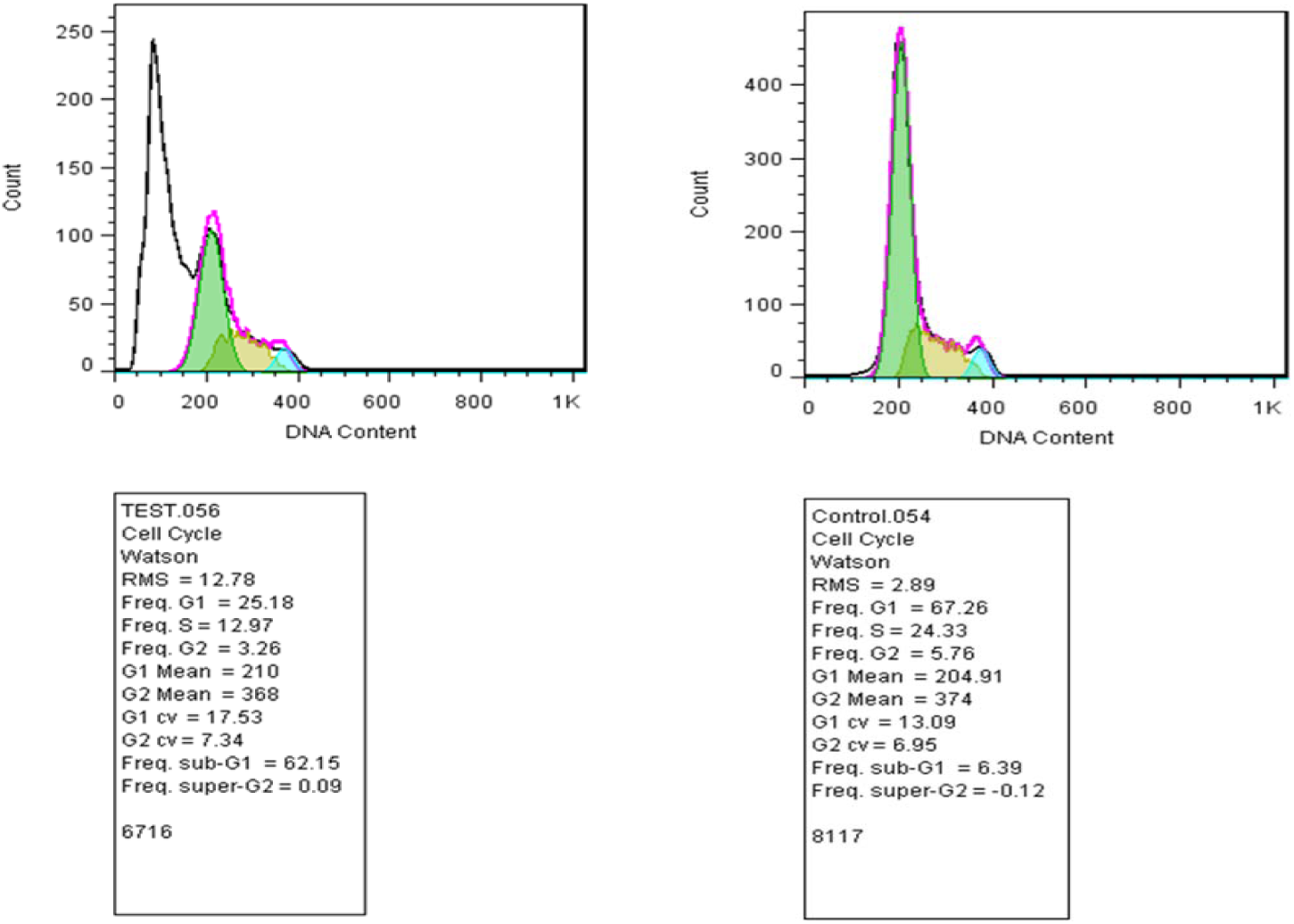
Representative flow cytometric analysis of cell cycle distribution in SKBR3 cells following treatment with 150 µM Silibinin. The left panel shows Silibinin-treated cells, and the right panel shows untreated control cells. DNA content histograms were analyzed to determine the percentage of cells in the G0/G1, S, and G2/M phases of the cell cycle. Silibinin treatment altered cell cycle progression compared with the untreated control.

**Figure 5** illustrates the percentage distribution of SKBR3 cells across the G1, G0/G1, S, and G2/M phases of the cell cycle. Consistent with the flow cytometry histograms shown in **Figure 4**, Silibinin treatment altered the cell cycle distribution of SKBR3 cells compared with the untreated control. Specifically, the proportion of cells in the G0/G1 phase decreased from 67.26% in the control group to 25.18% following Silibinin treatment. The percentage of cells in the G1 phase increased from 2.89% to 12.78%, whereas the S-phase population decreased from 24.33% to 12.97%. In addition, the proportion of cells in the G2/M phase decreased from 5.76% to 3.26%. Overall, these findings indicate that Silibinin significantly altered cell cycle progression in SKBR3 cells.

**Figure 5.**
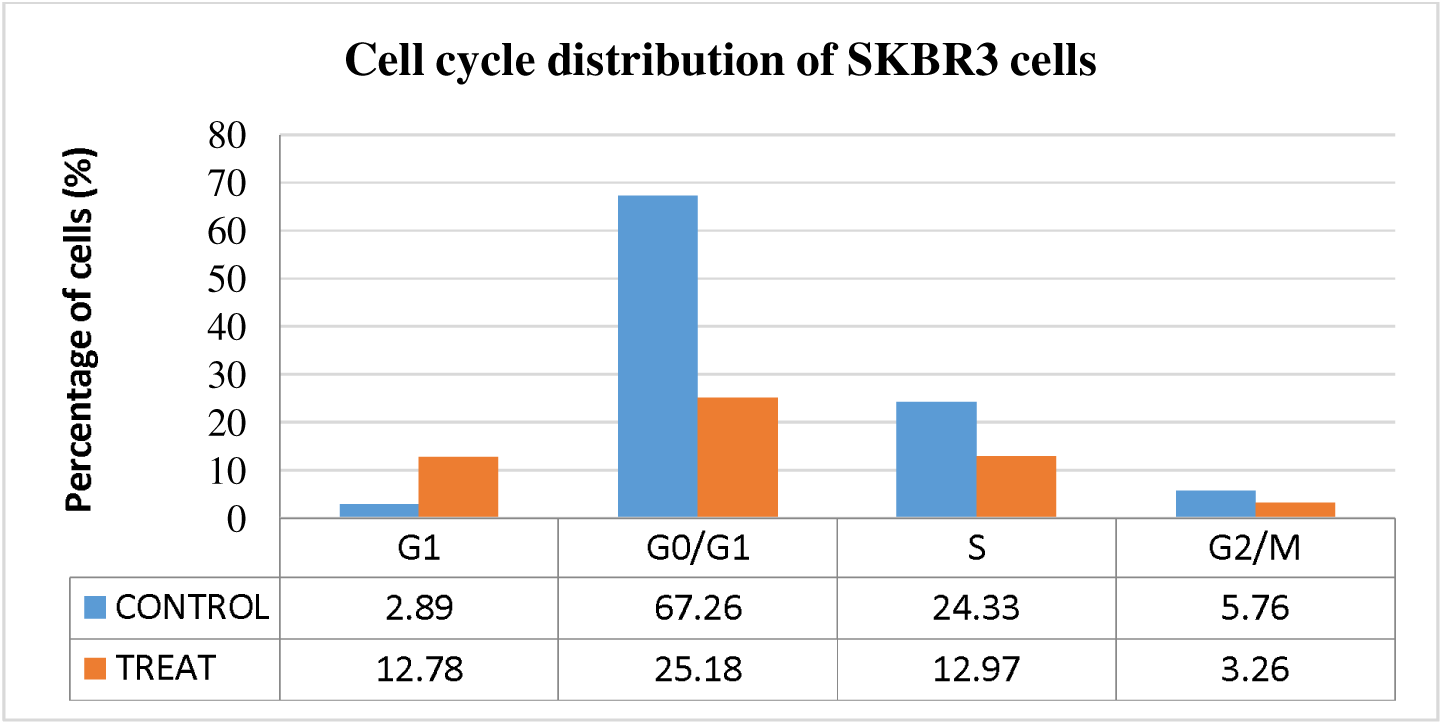
Cell cycle distribution of SKBR3 cells following treatment with 150 µM Silibinin. The percentages of cells in the G1, G0/G1, S, and G2/M phases were determined by flow cytometric analysis of DNA content. Data are presented as mean ± SD from three independent experiments (n = 3). Silibinin treatment altered the distribution of cells among the different phases of the cell cycle compared with the untreated control.

### ROS detection by DCFDA Assay

**Figure 6** shows the DSFDA assay results for ROS induction in the Silibinin-treated and untreated SKBR3 cells. According to the assay results, only 4% cells showed ROS production in the control group, as shown as ROS+. While, 94% ROS production was seen in the treatment sample, which is also shown as ROS+. Conversely, the percentage of ROSLJ cells (cells that do not release any fluorescent light) in the control sample was 96%, while the percentage of ROS + cells in the treatment sample was approximately 6%. According to **Figure 7**, the percentage of ROS-positive cells increased markedly from 4% in the control group to 94% in the Silibinin-treated group, while the percentage of ROS-negative cells correspondingly decreased from 96% to 6%, confirming that Silibinin treatment substantially elevated intracellular ROS levels in SKBR3 cells.

**Figure 6.**
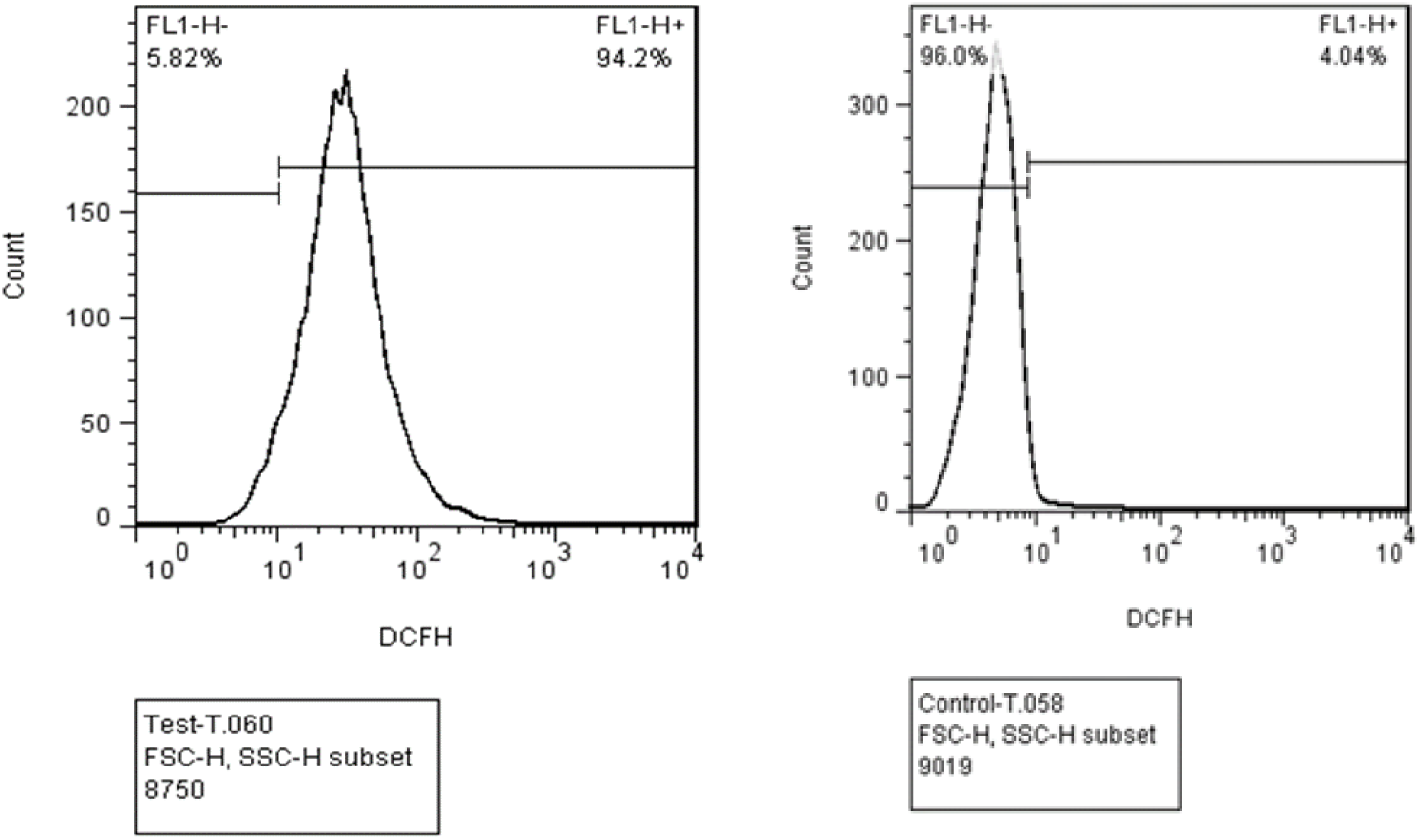
Representative flow cytometric histograms of intracellular reactive oxygen species (ROS) levels in SKBR3 cells following treatment with 150 µM Silibinin. Intracellular ROS generation was assessed using the DCFDA fluorescent probe. The left panel represents Silibinin-treated cells, whereas the right panel shows untreated control cells. A rightward shift in DCF fluorescence intensity indicates increased intracellular ROS production.

**Figure 7.**
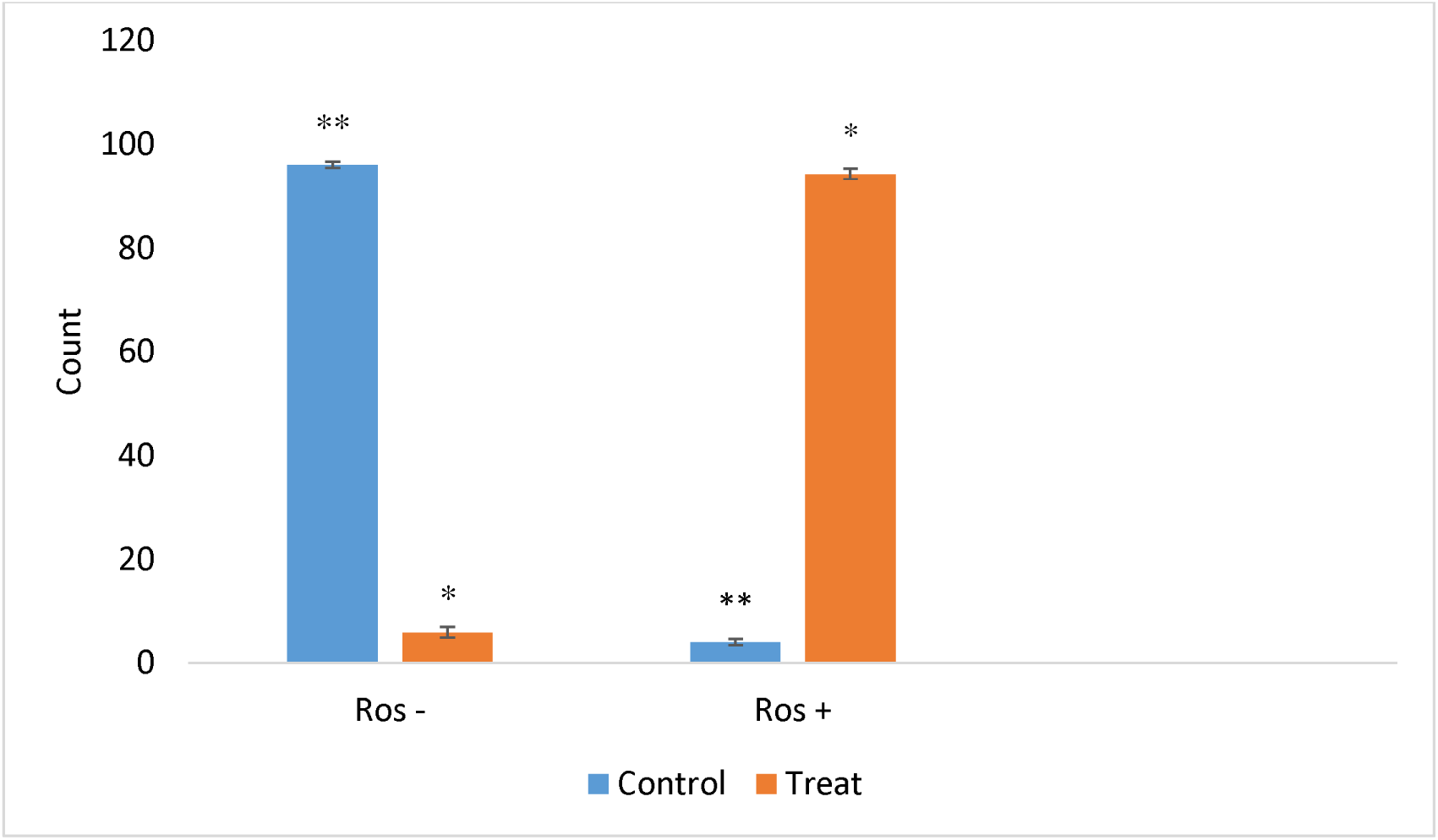
Comparison of intracellular reactive oxygen species (ROS) levels in untreated (control) and 150 µM Silibinin-treated SKBR3 cells, as determined by the DCFDA flow cytometric assay. ROS-negative (ROSLJ) and ROS-positive (ROSLJ) cell populations are presented as percentages of the total cell population. Data are presented as mean ± SD from three independent experiments (n = 3). Statistical significance was determined using one-way ANOVA followed by Tukey’s post hoc test. *P < 0.05 and **P < 0.01 versus the control group

### Real Time RT-PCR analysis of ATM gene expression in SKBR3 cells

**Figure 8** shows the RT-PCR analysis results in the real time. In this experiment, it was found that *ATM* gene expression in Silibinin-treated cells was higher than control samples, which shows that Silibininin could control *ATM* gene expression in SKBR3 cells.

**Figure 8.**
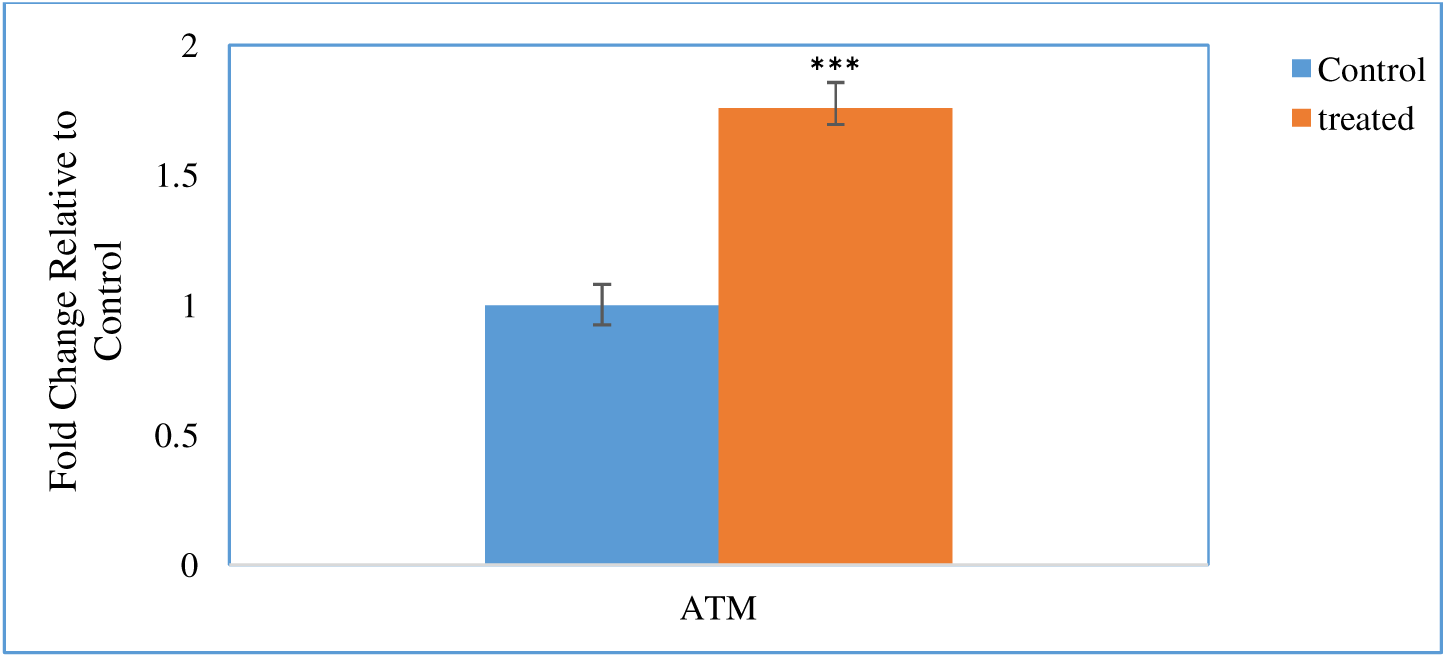
Relative ATM gene expression in untreated (control) and 150 µM Silibinin-treated SKBR3 cells as determined by quantitative real-time PCR (qRT-PCR). Gene expression levels were normalized to the internal reference gene and are presented as fold change relative to the untreated control. Data are expressed as mean ± SD from three independent experiments (n = 3). ***P < 0.001 versus the untreated control.

## Discussion

Recently, surgery, radiation therapy, chemotherapy, etc. are the conventional ways of breast cancer treatment. All these methods have disadvantages that have made the scientists find better ways to treat this disease [23]. Herbal medicine dates back several thousand years and the use of herbal products is one of better choices along with chemotherapeutic drugs, while reducing side effects of chemical drugs [24]. In this regards, the use of milk thistle in medical science has a long and brilliant history. More than 90% of the milk thistle extract comprises a string antioxidant phytochemical, Silibinin, which is considered to be the most effective ingredient in the seed extract of milk thistle. Moreover, no complications and side effects have been reported in human and animal models to date [25].

Many studies have reported the beneficial effects of the combination of milk thistle extract and chemotherapeutic drugs on breast, prostate, colon and skin cancers. Kim et al.(2016) showed that Silibinin inhibits TGF-β gene expression and metastatic initiation and progression in TNBC cells of breast cancer [26]. It has also been reported that Silibinin induces apoptosis in MCF-7 and MDA-MB-231 cells by increasing the ROS production in tumor microenvironment, which has also been proved by our results in SKBR3 cells [27]. This research group also found that the induction of apoptosis in MCF-7 cells is dependent on apoptosis-inducing factor (AIF), and in MDA-MB-231 cells, dependent on Caspase-3 protein [27].

The mechanism by which Silibinin increases the amount of ROS is not well defined. Several reports have shown that Silibinin can increase the ROS production through Ca^+2^ ion-dependent pathway [28]. Wang et al. (2010 and 2015) have reported that Silibinin increases the stability of extracellular superoxide production in the MCF-7 cell line [29]. In other studies on MCF-7 and T47D cell lines, it was found that Silibinin induces both internal and external apoptotic pathways in these cells, which is dependent on dose and incubation time [30, 31]. These results are in line with our results.

According to Pirouzpanah et al. (2015), Silibinin inhibits the growth of SKBR3 cells by inhibiting NF-βB. By studying the effect of Silibinin on apoptosis and cell cycle in MCF-7 cell line, these effects were reported due to increased expression of *ATM*, *p53*, *p21*, *Bak* and a decreased expression of *Bcl-xl* gene. All these cascades caused the cell cycle arrest at G0/G1 phase [32]. In a study by Wang et al. (2015), it was reported that silibillin causes impairment in mitochondrial function by producing ROS, which then resulted in autophagy in MCF-7 breast cancer cells [30]. In this study, it was found that by increasing the amount of ROS, Silibinin reduced the amount of ATP production in the mitochondria, which was effectively halted by adding antioxidant compounds, such as; N-acetylcysteine and ascorbic acid to the cell culture medium. This increase in ROS caused an increase in autophagy and cell death in MCF-7 cells [33]. Chavoshi et al. (2017) showed that the treatment of MCF-7 cells with a combination of paclitaxel, cisplatin and Silibinin could increase initial-stage apoptosis with an increase in *ATM*, *p53*, *Bax*, and *BRCA1* gene expressions [21].

More recent mechanistic work has reinforced and extended these findings. ATM is now recognized not only as a DNA-damage sensor but also as a direct redox sensor: oxidation of a cysteine pair within the ATM dimer promotes autophosphorylation and kinase activation independently of DNA double-strand breaks [35], a mechanism further resolved at the mitochondrial and structural level in subsequent studies [36,37]. This ROS-responsive branch of ATM signaling links oxidative stress directly to cell-cycle arrest, apoptosis and autophagy, and has been proposed as a therapeutic vulnerability in tumors with elevated oxidative burden [38]. Consistent with the present findings in SKBR3 cells, later studies on Silibinin in breast cancer have continued to implicate ROS-dependent pathways in its anticancer activity, including enhancement of immunogenic cell death and chemosensitivity [39], and disruption of Nrf2-dependent redox adaptation and JAK/STAT3 signaling in triple-negative breast cancer, further supporting a central role for oxidative stress and DNA-damage signaling in Silibinin’s anticancer mechanism across breast cancer subtypes [40].

## Conclusions

Taken together, the present study provides a coherent, multi-level picture of how Silibinin exerts its cytotoxic and pro-apoptotic effects on SKBR3 cells, a HER2-overexpressing human breast cancer cell line. At the cell-population level, MTT viability assays demonstrated that Silibinin reduced SKBR3 viability in both a dose- and time-dependent manner, with the corresponding IC50 decreasing from 234.49 µM at 24 h to 133.72 µM at 72 h, indicating that prolonged exposure progressively sensitized the cells to treatment. This cytotoxic effect was mechanistically linked, by flow cytometry, to a marked induction of apoptosis, evidenced by the shift of cells from the viable (Q4) to the early- and late-apoptotic (Q3 and Q2) quadrants of the Annexin V/PI assay, and to an accumulation of cells in the G1 phase of the cell cycle, consistent with a G1/S checkpoint arrest. At the molecular level, Silibinin treatment produced a substantial rise in intracellular ROS, from 4% ROS-positive cells in untreated controls to 94% in treated cells, together with a parallel increase in ATM gene expression. Since ATM functions both as a canonical DNA-damage sensor and, as more recent evidence indicates, as a direct redox sensor that is activated by oxidative modification independently of DNA double-strand breaks, these findings support a model in which Silibinin-induced ROS accumulation activates ATM signaling, which in turn engages the p53-dependent apoptotic pathway and enforces cell-cycle arrest at the G1/S transition. This sequence of events .oxidative stress, ATM/p53 activation, cell-cycle arrest and apoptosis — is consistent with the anticancer mechanisms reported for Silibinin in other breast cancer cell lines, including MCF-7, MDA-MB-231 and T47D, and extends these observations to the HER2-positive SKBR3 background, which had not been characterized with respect to ATM Signaling in this context. From a translational perspective, these results reinforce the rationale for evaluating Silibinin, either alone or in combination with conventional chemotherapeutic agents, as an adjunct strategy in HER2-positive breast cancer, where oxidative-stress- and DNA-damage-response pathways represent actionable vulnerabilities. Nevertheless, several limitations should be acknowledged. The present data derive from a single cell line and, for some endpoints, from a limited number of biological replicates; dose-response and time-course experiments should therefore be confirmed with additional HER2-positive and HER2-negative cell lines and with adequate statistical power before firm conclusions can be drawn regarding cell-line specificity. Likewise, the precise upstream mechanism linking Silibinin to ROS generation, and the causal relationship between ATM activation and the observed apoptotic and cell-cycle phenotypes, will require loss-of-function approaches (e.g., ATM knockdown or pharmacological inhibition) and direct measurement of ATM and downstream effector activation (such as p53, p21 and BAX) to be formally established. Future studies incorporating in vivo xenograft models and combination-treatment designs with standard-of-care chemotherapeutics would help clarify the therapeutic potential of Silibinin as a redox-and DNA-damage-response-targeting agent in HER2-positive breast cancer.

## Notes

### Competing Interest Statement

The authors have declared no competing interest.

